# Transient Splicing-Associated Isoform Remodeling Defines Early MAPKi Resistance Adaptation in Melanoma

**DOI:** 10.64898/2026.10.26.761932

**Authors:** Sumit Mukherjee, Arashdeep Singh, Sumeet Patiyal, Hyunjeong Joo, Hyungsoo Kim, Lipika R. Pal, Kun Wang, Chi-Ping Day, Ze’ev A Ronai, Eytan Ruppin, Sridhar Hannenhalli

## Abstract

A long-standing question in targeted therapy resistance is how tumor cells rapidly enter drug-tolerant states before stable resistance emerges. In BRAF-mutant melanoma, early adaptation to MAPK inhibitor (MAPKi) therapy is accompanied by profound phenotypic plasticity, yet the molecular mechanisms that enable this transition remain poorly understood. Here, we show that MAPKi adaptation is characterized by a transient transcript isoform remodeling that is largely invisible to conventional gene-expression analysis. We integrated and analyzed multiple different datasets, including bulk and single-cell RNA-seq datasets from melanoma cell lines, murine models, patient-derived xenografts and patient tumors, spanning no-treatment, on-treatment/minimal residual disease, and acquired-resistant states. We find that differential transcript usage peaks during the adaptive on-treatment phase and then is markedly attenuated when resistance was established. These dynamics were conserved across experimental systems and supported by independent exon-usage analyses. Notably, most genes undergoing differential transcript usage showed no corresponding change in total gene expression, and genes with isoform-level changes more strongly highlighted MAPKi resistance-associated pathways, including PI3K/AKT/mTOR and receptor tyrosine kinase signaling. Further, classifiers trained on cell-line transcript-usage features identified the adaptive state more accurately than gene-expression-based models in independent PDX and patient datasets. Interestingly, MAPKi-associated isoform changes recapitulated melanoblast-associated developmental splicing programs and were linked to a few key splicing regulators. Together, these findings identify an underappreciated regulatory layer connecting drug tolerance, developmental plasticity, and acquired resistance. Beyond the current context, our results suggest that comprehensively tracking isoform and splicing alterations may potentially advance our understanding of the emergence of treatment resistance in additional indications and treatment regimes.

## Introduction

Melanoma is one of the most aggressive forms of skin cancer that remains therapeutically challenging because of its strong propensity to develop treatment resistance (1). One of the major therapeutic advances for patients with BRAF-mutant melanoma has been the deployment of MAPK pathway inhibitors (MAPKi), which target the dysregulated MAPK signaling cascade driving tumor growth in a large fraction of melanomas. By inhibiting mutant BRAF and downstream MEK signaling, these targeted therapies can induce rapid tumor regression and improve clinical outcomes (2,3). However, responses are often incomplete and transient, and many patients ultimately develop resistance, limiting the long-term efficacy of MAPKi therapy (4,5).

Relapse is thought to arise from a small population of surviving melanoma cells that remain viable under drug pressure (6,7). These drug-tolerant persister cells can enter reversible adaptive states through non-mutational mechanisms before stable resistance is established (7,8). During this early on-treatment phase, residual surviving tumor cells constitute a minimal residual disease (MRD) state that can serve as a reservoir from which acquired resistance subsequently emerges. Previous studies have shown that adaptive resistance to MAPK inhibition is accompanied by heterogeneous transcriptional and phenotypic programs, including pigmentation, starvation, neural crest stem cell-like, and invasive states within residual disease (7,9). These observations support a model in which melanoma cells do not acquire resistance through a single linear trajectory but instead traverse dynamic drug-tolerant states that provide phenotypic plasticity during therapy adaptation.

Resistance to MAPKi is associated with diverse molecular alterations, including reactivation of MAPK signaling, activation of alternative survival pathways such as PI3K/AKT/mTOR signaling, receptor tyrosine kinase signaling, metabolic remodeling, and developmental cell-state reprogramming (6–10). Although gene-expression studies have provided important insights into these processes, they primarily quantify changes in total gene abundance and therefore may incompletely capture transcript-level changes occurring during adaptation (9). In particular, the contribution of transcript isoform remodeling to the early adaptive/MRD state remains less well characterized. Gene-level expression analysis can incompletely capture functionally relevant transcript changes when the overall expression of a gene remains unchanged, but the relative usage of its transcript isoforms is altered. Consequently, two cells with similar total expression of a gene may differ substantially in the relative abundance and functional properties of the transcripts produced from that gene. This distinction may be particularly important during early adaptation to MAPKi, when melanoma cells must rapidly and reversibly rewire signaling and cell-state programs to survive therapeutic pressure.

Differential transcript usage offers a complementary approach to define these adaptive transitions by capturing shifts in the relative abundance of transcript isoforms from the same gene. Such transcript-level changes can affect coding sequence, untranslated regions, regulatory elements, transcript stability, and pathway output, and many of these changes may arise through alternative splicing (11). Indeed, altered BRAF splicing and perturbation of pre-mRNA splicing have previously been implicated in MAPKi resistance in melanoma (12–14). However, whether dynamic transcript-usage changes broadly characterize the on-treatment state and provide information beyond differential gene expression remains unclear. Here, we integrated bulk and single-cell RNA-seq data from melanoma cell lines, patient tumors, murine melanoma models, and patient-derived xenografts spanning no-treatment, on-treatment/MRD, and acquired-resistant states. We found that MAPK inhibition induces a pronounced but transient rewiring of transcript usage that peaks during the on-treatment adaptive/MRD phase and is largely attenuated after resistance is established. Although the specific transcripts altered across melanoma models were not always identical, treatment-associated isoform changes showed conserved temporal and pathway-level dynamics. These transient transcript changes were also supported by independent exon-usage analyses. Importantly, most genes with altered transcript usage did not show corresponding changes in total gene expression, indicating that transcript-level remodeling captures an adaptive layer that is not fully resolved by differential gene-expression analysis. Consistent with this, isoform-level features showed stronger enrichment for MAPKi resistance-associated pathways and more accurately identified the adaptive on-treatment state in independent melanoma datasets than gene-expression features. Finally, integration with splicing-factor and developmental transcriptomic analyses linked this transient isoform program to melanoblast development-associated cellular plasticity. Together, these findings identify transient transcript isoform remodeling as a defining feature of early MAPKi resistance adaptation and demonstrate that isoform-level analysis provides a more informative molecular readout of the adaptive/MRD state than gene-expression analysis alone.

## Results

### 1. Overview of the analysis and datasets

We designed a multi-step computational framework to investigate transcript isoform remodeling during the development of MAPKi resistance in melanoma and to determine whether isoform-level changes capture adaptive responses beyond conventional gene-expression analysis **(Figure 1)**. First, we analyzed longitudinal transcriptomic data from MAPKi-treated melanoma cell lines spanning no-treatment, on-treatment/adaptive, and acquired-resistant states, using the GSE75299 dataset as the primary discovery cohort (6). Transcript-level quantification and differential transcript usage analyses were performed to identify dynamic isoform changes across treatment states **(Figure 1a)**. We then evaluated the reproducibility of these treatment-associated patterns across complementary melanoma models, including patient samples from GSE75299, an independent murine melanoma dataset (GSE103725) (6), and the GSE116237 single-cell RNA-seq PDX dataset (7), for which pseudo-bulk transcriptomes were generated to enable transcript-level analysis across sequential treatment phases **(Figure 1a,b)**. The PDX dataset includes pre-treatment and multiple MAPKi treatment phases extending to resistance.

**Figure 1.**
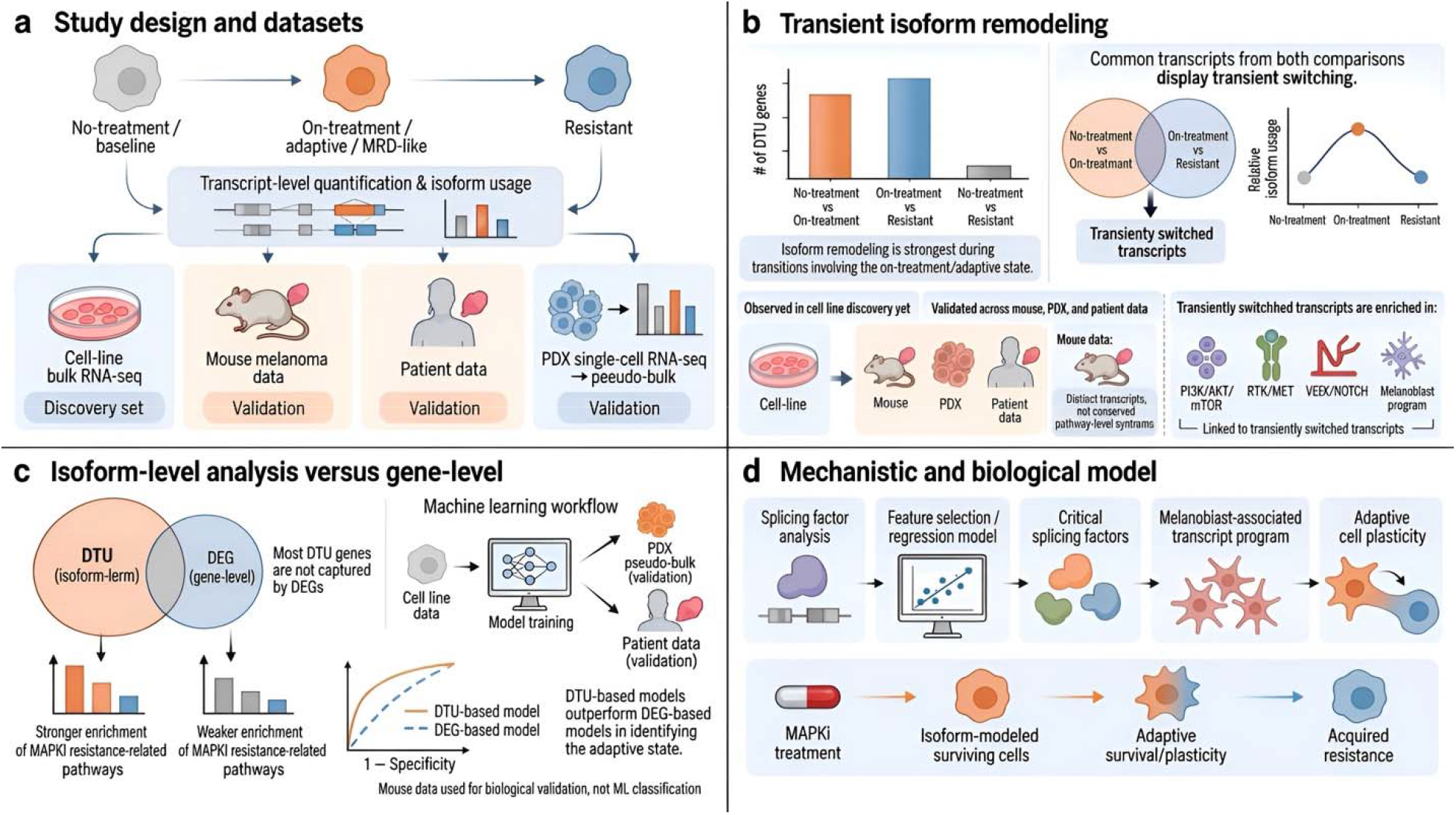
Study design and analytical framework for characterizing transcript isoform remodeling during MAPKi resistance evolution in melanoma. **(a)** Overview of the transcriptomic datasets and treatment states analyzed in this study. Melanoma cell-line bulk RNA-seq spanning no-treatment/baseline, on-treatment/adaptive or MRD-like, and acquired-resistant states was used as the primary discovery dataset. Transcript-level quantification and isoform-usage analysis were performed across these treatment states, and the resulting patterns were subsequently evaluated in complementary validation datasets, including murine melanoma, patient samples, and PDX single-cell RNA-seq data summarized as pseudo-bulk transcriptomes. **(b)** MAPKi-associated isoform remodeling was most prominent during transitions involving the on-treatment/adaptive state. Transcripts common to the no-treatment versus on-treatment and on-treatment versus resistant comparisons exhibited transient switching, with isoform usage changing during treatment and subsequently reversing as resistance emerged. These treatment-associated dynamics were first identified in the cell-line discovery dataset and then examined across mouse, PDX, and patient data. Although the specific transcripts detected in the murine dataset were not necessarily identical to those observed in the human cell-line data, they showed comparable temporal behavior and convergence on similar pathway-level programs. Transiently switched transcripts were enriched in biological processes associated with MAPKi adaptation and resistance, including PI3K/AKT/mTOR, RTK/MET, VEGF/NOTCH, and melanoblast-associated programs. **(c)** Comparison of isoform-level differential transcript usage (DTU) with conventional gene-level differential expression (DEG). Most genes exhibiting significant DTU were not captured by DEG analysis, indicating that isoform remodeling represents an additional layer of treatment-associated regulation. DTU genes also showed stronger enrichment for MAPKi resistance-related pathways than DEGs. To assess the predictive information contained at each molecular level, machine-learning models were trained using the cell-line discovery data and independently evaluated in PDX pseudo-bulk and patient datasets. Isoform-level models provided a more informative readout of the adaptive on-treatment state than gene-level models. Mouse data were used for biological validation of conserved treatment-associated programs and were not included in machine-learning classification. **(d)** Mechanistic and biological model of MAPKi-associated isoform remodeling. Splicing-factor expression was integrated with feature selection and regression modeling to identify critical splicing regulators associated with the transient transcript program. These regulatory patterns were further linked to melanoblast-associated transcript usage and adaptive cell plasticity. Together, the analyses support a model in which MAPKi treatment induces a transient isoform-remodeled state in surviving melanoma cells, enabling adaptive survival and phenotypic plasticity before progression toward acquired resistance.

Second, we compared differential transcript usage with gene-level differential expression to determine whether isoform remodeling provides additional information about MAPKi adaptation. Models derived from the cell-line data were subsequently evaluated in independent PDX pseudo-bulk and patient data to compare the ability of isoform- and gene-level features to correctly classify the adaptive on-treatment state **(Figure 1c)**. Finally, we investigated potential regulators and biological consequences of the transient isoform program by integrating splicing-factor modeling with developmental transcriptomic analyses, linking MAPKi-associated isoform remodeling to melanoblast development-associated programs and adaptive cellular plasticity **(Figure 1d)**. Together, this framework enabled us to characterize the temporal dynamics, cross-model conservation, predictive relevance, and biological basis of transcript isoform remodeling during MAPKi resistance evolution in melanoma.

### 2. MAPKi treatment induces a conserved transient transcript-usage program in on-treatment melanoma cells

To determine whether MAPKi treatment reshapes transcript usage during the evolution of acquired resistance, we first analyzed bulk RNA-seq profiles from MAPKi-treated melanoma cell lines in the GSE75299 dataset, which profiled MAPKi-induced melanoma regression and early adaptation. This dataset includes melanoma cell lines sampled across three states: untreated baseline, on-treatment drug-tolerant/MRD-like cells, and fully resistant cells. Differential transcript usage (DTU) analysis was performed across three pairwise comparisons: no-treatment versus on-treatment, on-treatment versus resistant, and no-treatment versus resistant. The largest transcript-usage alterations were observed during transitions involving the on-treatment state. While only 147 genes showed DTU between no-treatment and resistant cells, 1,099 genes showed significant DTU between no-treatment and on-treatment cells, and 1,569 genes showed significant DTU between on-treatment and resistant cells. This pattern indicates that transcript-usage remodeling is most prominent during the adaptive on-treatment phase rather than in the final resistant state **(Figure 2a)**.

**Figure 2.**
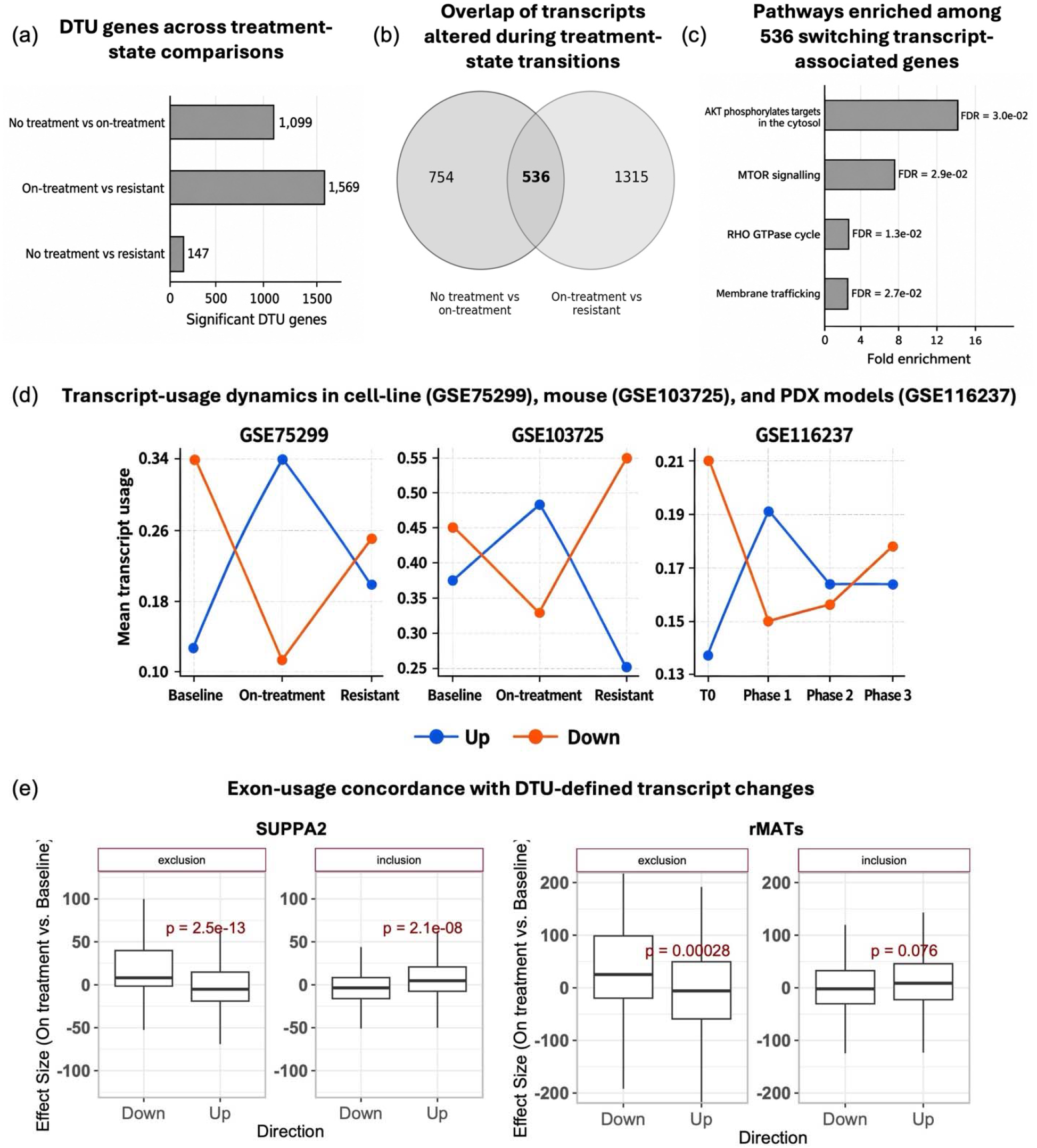
MAPKi treatment induces a conserved transient transcript-usage program across melanoma models. **(a)** Number of genes with significant differential transcript usage (DTU) across treatment-state comparisons in MAPKi-treated melanoma cell lines (GSE75299). DTU alterations were markedly more abundant in comparisons involving the on-treatment state, with 1,099 DTU genes identified between no-treatment and on-treatment cells and 1,569 between on-treatment and resistant cells, compared with only 147 between no-treatment and resistant cells. **(b)** Overlap of transcripts altered during the no-treatment versus on-treatment and on-treatment versus resistant transitions in GSE75299 cell line dataset. A total of 536 transcripts were altered in both transitions, consistent with a transient switching pattern during MAPKi adaptation. **(c)** Pathway overrepresentation analysis of genes associated with the 536 switching transcripts, highlighting enrichment of AKT cytosolic phosphorylation targets, mTOR signaling, RHO GTPase cycle, and membrane trafficking. **(d)** Mean transcript-usage dynamics of transcripts increased or decreased during the on-treatment state across melanoma cell-line (GSE75299), murine melanoma (GSE103725), and PDX models (GSE116237). In the cell-line discovery dataset (GSE75299), transcripts increased during treatment showed a peak in the on-treatment state and subsequently declined in resistant cells, whereas transcripts decreased during treatment showed the reciprocal pattern. A similar transient trajectory was observed in the independent murine melanoma dataset (GSE103725), despite differences in the specific transcripts involved. In the PDX pseudo-bulk dataset (GSE116237), concordant changes were most evident in the phase 1 on-treatment/MRD state and were attenuated during later treatment phases, supporting conservation of the transient adaptive isoform program across experimental systems. **(e)** Independent exon-usage validation of DTU-defined transcript changes in the GSE75299 MAPKi-treated melanoma cell-line dataset using SUPPA2 and rMATS. Exons corresponding to transcripts with increased usage during treatment showed concordant increases in exon inclusion, whereas exons associated with transcripts showing decreased usage exhibited the opposite pattern, supporting agreement between transcript-level DTU and exon-level splicing analyses.

We next identified transcripts that changed direction across the resistance trajectory. This analysis revealed 536 transcripts whose usage changed significantly between no-treatment and on-treatment cells and then reversed from on-treatment to resistant cells and consequently were not detected as differentially used in the direct comparison of no-treatment versus resistant cells **(Figure 2b)**. These results indicate that a substantial fraction of transcript-usage alterations is transient, emerging under MAPKi exposure and then attenuating after resistance is established. Pathway enrichment analysis of genes from these 536 switching transcripts revealed AKT phosphorylates targets in cytosol and mTOR signaling among the top pathways **(Figure 2c)**. These findings link transient transcript switching to PI3K/AKT/mTOR-associated signaling, RHO GTPase regulation, and membrane-trafficking processes during MAPKi adaptation. The transient transcript-usage dynamics of these 536 transcripts across the three treatment states are depicted in **Figure 2d** (*left panel*).

To evaluate whether the early MAPKi-induced DTU patterns detected in cell lines were reproduced in additional melanoma models, we analyzed an independent murine melanoma dataset (GSE103725). Because the murine dataset represents a different species, the specific transcripts identified in the human GSE75299 cell-line dataset could not be directly mapped on a one-to-one basis. Instead, we asked whether the overall transcript-usage dynamics were conserved across treatment states. In the murine model, transcripts showing increased or decreased usage during the on-treatment state displayed the same transient pattern observed in the human cell-line data, with reciprocal changes across baseline, on-treatment, and resistant states **(Figure 2d**, *middle panel*). Further, pathway enrichment analysis of genes exhibiting transient transcript-usage dynamics in the murine model revealed significant enrichment of PI3K/AKT signaling (FDR = 0.036) and ErbB signaling (FDR = 0.028), both implicated in adaptive MAPKi resistance. Notably, PI3K/AKT signaling was also enriched in the human cell-line analysis, supporting convergence of resistance-associated signaling programs across models. These findings suggest that, despite differences in individual transcripts, transient isoform remodeling involves shared MAPKi resistance-associated pathways in human and murine melanoma.

Next, we analyzed a patient-derived xenograft model profiled by single-cell RNA-seq across treatment phases from GSE116237. For this dataset, pseudo-bulk transcriptomes were generated to reduce the impact of single-cell dropout. Transcripts identified as increased or decreased during the baseline-to-on-treatment transition in GSE75299 cell-line dataset showed concordant changes in phase 1 on-treatment/MRD PDX samples, with these patterns attenuating during later treatment phases **(Figure 2d**, *right panel*). Together, these findings show that MAPKi induces a transient transcript-usage program that is reproducible across melanoma models, most evident in on-treatment/MRD melanoma cells, and only partially persists in fully resistant tumors.

To further support the DTU findings, we additionally quantified Percent-Spliced-In (PSI) values of alternative exon usage events associated with DTU events between baseline and on-treatment conditions (Methods). We observed that exons included in transiently upregulated transcripts were also upregulated in exon-usage analyses, whereas exons excluded from transiently upregulated transcripts showed reduced usage **(Figure 2e)**. Thus, independent exon-level analyses were broadly concordant with the transcript-level DTU results, supporting the robustness of the detected treatment-associated splicing-associated transcript remodeling.

### 3. Transcript-level remodeling captures additional layers of adaptive MAPKi-response programs not fully resolved at the gene-expression level

We next asked whether MAPKi-associated transcript-usage alterations could be explained by changes in total gene expression. To address this, we compared DTU genes detected in each treatment contrast with differentially expressed genes (DEGs) identified from the corresponding gene-level comparison using DESeq2. In both the no-treatment versus on-treatment and on-treatment versus resistant comparisons, most genes with significant transcript-usage alterations did not show corresponding changes in total gene expression **(Figure 3(a))**. Only a small fraction (11.5% in the no-treatment versus on-treatment comparison) of genes undergoing transcript-usage changes overlapped with DEGs, indicating that MAPKi-induced transcript remodeling is largely uncoupled from gene-level expression changes. These findings support a model in which melanoma cells under MAPKi pressure undergo isoform-level remodeling that is not captured by total gene expression. Such changes may allow cells to alter transcript architecture and pathway output while maintaining stable gene-level expression. Thus, transcript-usage analysis reveals an additional layer of treatment-associated adaptation in on-treatment and MRD-like melanoma cells.

**Figure 3.**
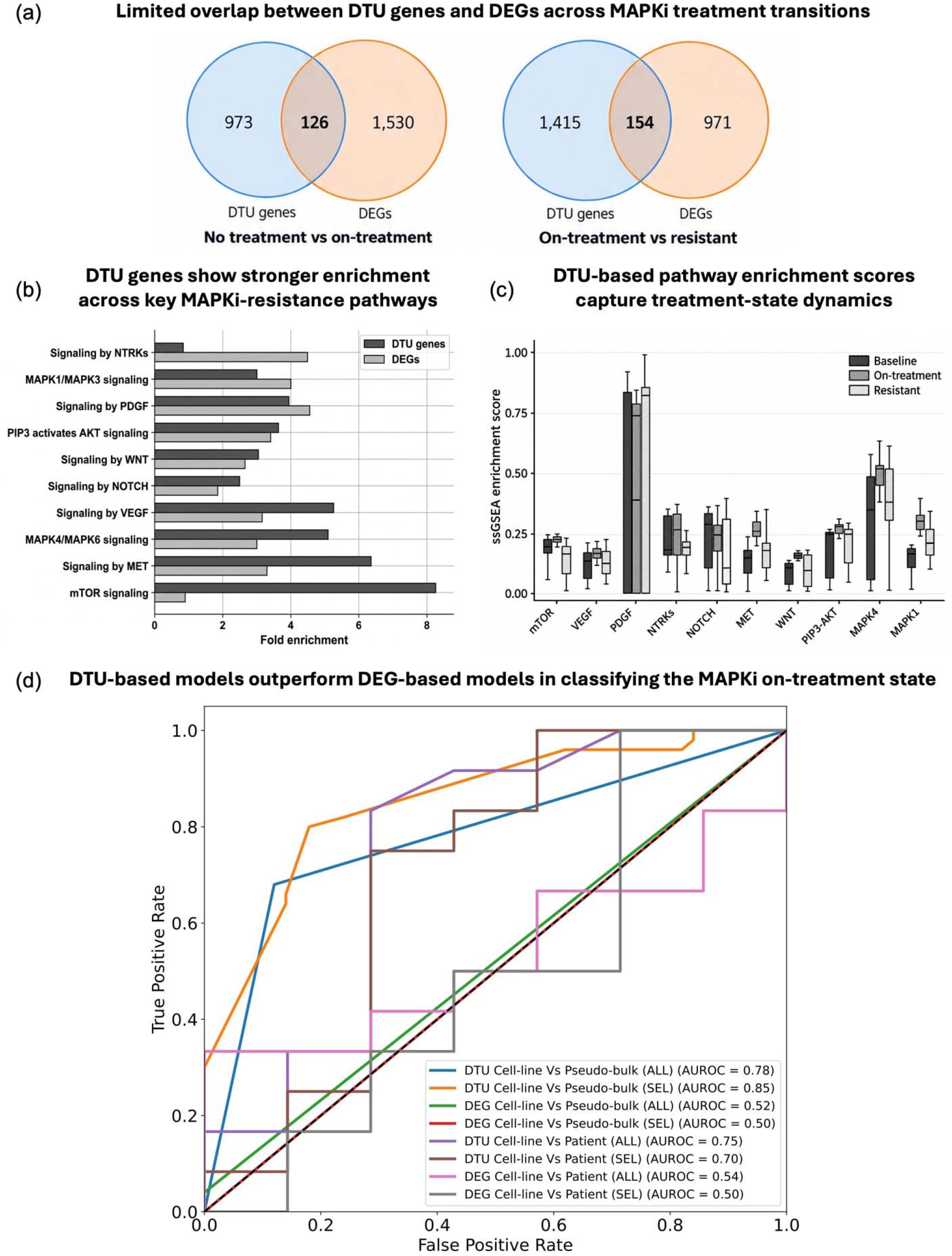
Transcript-usage remodeling is largely uncoupled from gene expression and predicts the MAPKi-adaptive state. **(a)** Overlap between DTU genes and differentially expressed genes (DEGs) in the no-treatment versus on-treatment and on-treatment versus resistant comparisons. In both treatment-state transitions, only a small fraction of DTU genes overlapped with DEGs, indicating that MAPKi-associated transcript-usage remodeling is largely uncoupled from changes in total gene expression. **(b)** Pathway overrepresentation analysis comparing DTU genes and DEGs detected in the no-treatment versus on-treatment comparison in the GSE75299 cell-line dataset. Fold enrichment is shown for selected MAPKi resistance-associated pathways, including receptor tyrosine kinase signaling, PI3K/AKT signaling, MAPK signaling, WNT, NOTCH, MET, VEGF, PDGF, and mTOR pathways. DTU genes showed stronger enrichment across several resistance-associated pathways, supporting transcript-level remodeling as an additional readout of adaptive MAPKi-response biology. **(c)** ssGSEA enrichment scores for transcripts with increased usage in on-treatment cells across baseline, on-treatment, and resistant states in the GSE75299 cell-line dataset. Transcript-usage–derived pathway scores varied across treatment states, indicating that DTU captures dynamic modulation of resistance-associated signaling programs during MAPKi response and resistance evolution. **(d)** Receiver operating characteristic analysis comparing DTU- and DEG-based models. DTU-based classifiers outperformed DEG-based classifiers across testing conditions, with the selected-feature DTU model achieving the highest performance in pseudo-bulk PDX data, whereas DEG-based models performed close to random. These analyses show that transcript-usage features provide a more predictive readout of the MAPKi-adaptive on-treatment state than gene-expression features.

To determine whether these gene-expression-independent transcript-usage changes captured resistance-relevant biology, we next compared enriched pathway between DTU genes and DEGs detected in the no-treatment versus on-treatment comparison in the GSE75299 cell-line dataset. We focused on key pathways previously implicated in MAPKi resistance (15–21). Although both DTU genes and DEGs were enriched for several resistance-associated pathways, DTU genes showed stronger enrichment across most of the examined pathways **(Figure 3b)**. The difference was particularly evident for mTOR, MET, MAPK4/MAPK6, VEGF, and NOTCH signaling, indicating that transcript-level remodeling captures pathway alterations that are less apparent from gene-level expression changes alone. To further examine pathway modulation across treatment states in the GSE75299 cell-line dataset, we performed ssGSEA using transcripts with increased usage in on-treatment cells. This analysis showed dynamic enrichment-score patterns across baseline, on-treatment, and resistant samples for several MAPKi resistance-associated pathways **(Figure 3c)**. These results indicate that transcript-usage changes are linked to treatment-state-dependent shifts in resistance-associated signaling programs. Together, the overrepresentation and ssGSEA analyses further support the findings that DTU provides a more informative readout of adaptive MAPKi-response pathways than DEG analysis.

We next directly tested whether DTU or DEG features better classify the adaptive on-treatment state. Machine-learning classifiers were trained using cell-line bulk RNA-seq data to distinguish pre-treatment from on-treatment samples. To assess generalizability of the models, the cell-line-trained models were independently evaluated in both pseudo-bulk transcriptomes generated from the PDX single-cell dataset and an independent patient dataset **(Figure 3d)**. Models were evaluated using all detected features as well as selected transcript or gene features (*see methods*). In the PDX pseudo-bulk validation, DTU-based models clearly outperformed DEG-based models: the DTU model achieved an AUC of 0.78 using all features and 0.85 after feature selection, whereas the corresponding DEG-based models achieved AUCs of 0.52 and 0.50, respectively. A similar advantage was observed in the independent patient dataset, where DTU-based models achieved AUCs of 0.75 using all features and 0.70 after feature selection, compared with 0.54 and 0.50 for the corresponding DEG-based models. Thus, despite differences in model system and sample composition, transcript-usage features learned from the cell-line data retained substantially greater discriminatory ability than gene-level features in both independent validation settings. Together, these results show that DTU better captures the adaptive transcriptional programs associated with MAPKi exposure than DEG. Thus, transcript-usage analysis reveals MAPKi-resistance programs that are incompletely captured by gene-level expression analysis and are more predictive of on-treatment samples in independent cohorts.

### 4. MAPKi-associated isoform remodeling recapitulates melanoblast developmental programs and is linked to splicing-factor regulation

Past research has shown that melanoma cells, following MAPKi treatment, can undergo phenotypic switching to a slow-cycling, melanoblast-like differentiation state via MITF driven transcriptional reprogramming (7,22). Furthermore, in multiple cancer types the global splicing program in the tumor has been shown to recapitulate the corresponding developmental splicing program (23). Therefore, we investigated if the alternative splicing events that characterize the melanocytic precursor phenotype are re-activated in on-treatment drug-tolerant cells. Towards this, we analyzed an independent dataset (24) consisting of *in-vitro* melanocytic differentiation from neural crest-like cells and compared melanoblasts and mature melanocytes. We processed the RNA-seq data from the WT models (*See Methods*) and identified transcript-usage events differentially regulated during melanoblast-to-melanocyte transition at an FDR threshold of 0.05. As a sanity check, we performed the functional enrichment analysis of the corresponding genes of the transcripts that showed greater inclusion levels in the precursor melanoblast relative to differentiated melanocytic state and observed that top enriched function such as cytoskeleton and actin filament organization, were consistent with the spindle shaped morphology as well migratory behavior of the melanoblasts (**Supplementary Figure 1a**). Next, we tested whether transient MAPKi-induced transcript changes recapitulated the melanoblast-associated developmental splicing signature. We observed that transcripts upregulated during the on-treatment state showed significantly higher melanoblast-versus-melanocyte delta PSI values than downregulated transcripts, indicating that MAPKi-induced transcript remodeling is biased toward melanoblast-associated transcript usage **(Figure 4a)**. Further, using a more stringent set of melanoblast specific transcripts (FDR < 0.05 & > 10% change in inclusion level during melanocyte differentiation) further strengthened this trend (**Supplementary Figure 1b)**. In particular, we observed that ROBO2 and SRCIN1 genes, which are critical for the migratory behavior of melanoblasts have alternatively spliced isoforms which were re-activated upon MAPKi treatment in melanoma cell lines (**Figure 4a and Supplementary Figure 1c**)

**Figure 4.**
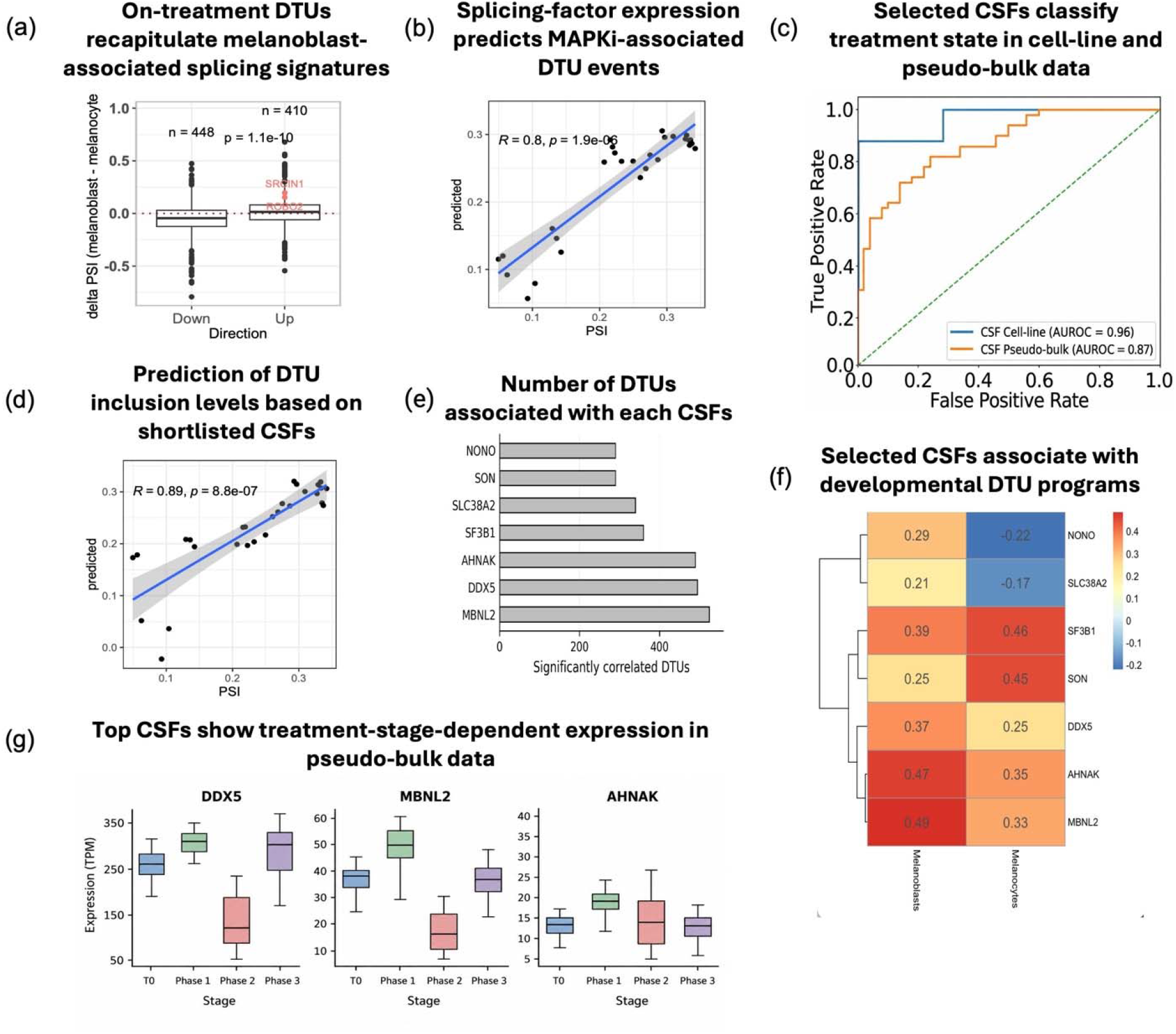
Critical splicing factors link transient MAPKi-induced transcript remodeling to developmental melanoblast-associated programs. **(a)** Recapitulation of developmental splicing signatures in on-treatment melanoma cells. Transcripts upregulated during the on-treatment state showed significantly higher melanoblast-versus-melanocyte delta PSI values than downregulated transcripts, indicating enrichment of melanoblast-associated transcript usage during MAPKi adaptation. **(b)** Partial least squares regression (PLSR) model using expression profiles of annotated splicing factors to predict MAPKi-associated DTU inclusion levels. The model showed strong agreement between observed and predicted inclusion levels (R = 0.80, *P = 1.9 × 10*⁻⁶), supporting an association between splicing-factor expression and MAPKi-induced transcript-usage remodeling. **(c)** Receiver operating characteristic analysis evaluating the selected critical splicing-factor (CSF) model for treatment-state classification. The selected CSF model showed strong performance in both cell-line data (AUROC = 0.96) and pseudo-bulk PDX data (AUROC = 0.87). **(d)** PLSR evaluation after machine-learning-based feature selection. The selected CSF model improved prediction of observed DTU inclusion levels (R = 0.89, P = *8.8 × 10*⁻⁷) and prioritized seven CSFs: MBNL2, AHNAK, NONO, DDX5, SF3B1, SLC38A2, and SON. **(e)** Number of DTU events significantly associated with each of the seven selected CSFs. MBNL2, DDX5, and AHNAK were associated with the largest numbers of DTU events, highlighting their potential involvement in MAPKi-induced transcript-usage remodeling. **(f)** Association of selected CSFs with DTUs enriched in melanoblast- and melanocyte-associated developmental programs. The selected CSFs showed distinct developmental associations, with the leading MAPKi-associated CSFs showing positive association with melanoblast-linked DTU programs. **(g)** Expression dynamics of the top three CSFs across pseudo-bulk PDX treatment stages. DDX5, MBNL2, and AHNAK showed treatment-stage-dependent expression patterns, with increased expression during the phase 1 on-treatment state relative to T0.

Having established the association between MAPKi-induced transcript remodeling and melanoblast developmental programs, we next investigated whether the observed DTU events were associated with coordinated changes in splicing-factor expression. To this end, we curated 442 annotated splicing factors and employed partial least squares regression (PLSR) to assess their association with the transient transcript-usage program. Specifically, the expression levels of the 442 splicing factors were used as predictor variables, while the mean inclusion level of transiently upregulated transcripts (TUTs) across samples served as the response variable. This approach allowed us to examine whether splicing-factor expression could explain variation in the transcript-usage program associated with MAPKi treatment. We identified 85 splicing factors with positive regression coefficients and statistically significant associations (FDR < 0.05). The resulting PLSR model showed a strong correlation between observed and predicted transcript inclusion levels (R = 0.80 and P = *1.9 × 10*) **(Figure 4b)**.

We then refined this candidate set further using machine-learning-based feature selection followed by PLSR evaluation. The selected-factor classifier showed strong performance in distinguishing treatment states, with an AUROC of 0.96 in cell-line data and 0.87 in pseudo-bulk PDX data **(Figure 4c)**. This analysis prioritized seven splicing regulators: MBNL2, AHNAK, NONO, DDX5, SF3B1, SLC38A2, and SON. Using these seven factors, the reduced PLSR model showed improved concordance between observed and predicted transcript inclusion levels (R = 0.89 and *P = 8.8 × 10*) **(Figure 4d)**. Together, these analyses indicate that these seven splicing-factors are sufficient to capture MAPKi-associated transcript-usage remodeling and nominate these splicing factors as critical splicing factors (CSFs). Among these CSFs, MBNL2, DDX5, and AHNAK were associated with the largest number of significantly correlated DTUs **(Figure 4e)** as well as showed preferential expression in melanoblast precursors as compared to differentiated melanocytes (**Figure 4f**). We therefore further examined their expression dynamics in pseudo-bulk PDX data across treatment stages. All these three CSFs showed treatment-stage-dependent expression patterns, with increased expression during the phase 1 on-treatment state relative to T0 **(Figure 4g)**. These results prioritize MBNL2, DDX5, and AHNAK as leading splicing-factor candidates associated with the transient MAPKi-induced transcript-usage state. Together, these analyses support a model in which MAPKi treatment induces a transient transcript-usage state associated with a critical splicing-factor module, led by MBNL2, DDX5, and AHNAK, and linked to reactivation of melanoblast-associated developmental splicing programs.

## Discussion

Acquired resistance to MAPKi remains a central limitation in the treatment of BRAF-mutant melanoma. Our study identifies transient splicing-associated transcript-usage remodeling as a previously underappreciated feature of the adaptive response to MAPKi. Rather than being most prominent in the fully resistant state, transcript-usage changes were strongest during the on-treatment/MRD-like phase and were largely attenuated after resistance was established. This temporal behavior places isoform remodeling within the early adaptive window that precedes stable resistance, rather than as a fixed molecular consequence of the resistant state itself. This pattern suggests that transcript-level remodeling reflects a dynamic adaptive program that emerges under therapeutic pressure and may contribute to the phenotypic flexibility required for survival during minimal residual disease. Importantly, similar transient dynamics were observed across cell-line, murine melanoma, and PDX models, indicating that although the specific transcripts affected can differ between systems, the broader temporal program is conserved.

A key finding of this work is that transcript-usage remodeling is largely uncoupled from total gene-expression changes during early adaptation to therapeutic pressure. Most genes with altered transcript usage did not show corresponding changes in gene-level expression, indicating that melanoma cells can remodel transcript architecture without substantially altering overall gene abundance. This distinction is important because conventional differential gene-expression analysis may miss adaptive programs that operate through isoform switching, altered transcript composition, or changes in regulatory regions. Thus, gene-level expression and transcript-usage analyses capture partially distinct layers of the adaptive response. Consistent with this, DTU-based classifiers distinguished the pre-treatment and on-treatment states more effectively than DEG-based classifiers, including in independent PDX pseudo-bulk and patient validation datasets. The reproducibility of this advantage across distinct validation settings suggests that isoform-level information captures a generalizable feature of the MAPKi-adaptive state rather than a cohort-specific signal. These results support the concept that transcript-usage features provide a sensitive readout of the MAPKi-adaptive state.

Transcript-level analysis also provided stronger functional insight into pathways associated with resistance evolution. DTU genes more prominently highlighted PI3K/AKT signaling, receptor tyrosine kinase signaling, mTOR-associated programs, cellular signaling and trafficking, and lipid metabolic remodeling. The enrichment of transient switching transcripts in mTOR signaling, RHO GTPase regulation, and membrane-trafficking processes further connects transcript-usage dynamics with signaling and cellular adaptations accompanying MAPKi exposure. These pathway-level findings are notable because they connect the isoform changes to signaling networks already implicated in adaptive MAPKi resistance, suggesting that transcript remodeling may modulate known resistance pathways at a regulatory level not readily visible from total gene abundance. Rather than replacing gene-expression changes, isoform remodeling may therefore complement transcriptional adaptation by altering transcript composition and potentially modifying pathway output during the drug-tolerant phase. These findings suggest that transcript-level remodeling may help melanoma cells rapidly modulate pathway output during the drug-tolerant phase, thereby supporting a permissive state for survival under MAPKi pressure.

Our analysis also connects MAPKi-induced transcript remodeling with developmental plasticity. The enrichment of melanoblast-associated transcript-usage patterns in on-treatment cells suggests that MAPKi exposure promotes a partial reactivation of developmental programs during adaptation. This is consistent with the broader model that melanoma cells can access neural crest-like or progenitor-like states under therapeutic stress (7,25,26). In this context, transient transcript-usage rewiring may provide an additional post-transcriptional mechanism that complements genetic, epigenetic, and transcriptional programs through which melanoma cells acquire phenotypic flexibility during therapy adaptation (27,28). Further, the splicing-factor analysis provides a potential regulatory link between MAPKi exposure and this transient isoform program. Our modeling prioritized a set of critical splicing factors, including MBNL2, DDX5, and AHNAK, that were associated with MAPKi-induced transcript-usage changes and showed treatment-stage-dependent expression patterns. These factors were also preferentially associated with melanoblast-related transcript programs, suggesting that coordinated changes in splicing-factor activity may contribute to the developmental plasticity observed during early adaptation. However, these analyses are correlative and should be interpreted as nominating candidate regulators rather than establishing direct causal control of the identified isoforms.

Several limitations should be noted. First, this study is based primarily on publicly available RNA-seq datasets, and the timing, treatment regimens, sequencing platforms, and model systems differ across cohorts. Although this heterogeneity provides an opportunity to test robustness and reproducibility of the signals, it also limits direct quantitative comparison of individual transcript changes across datasets. Second, although independent exon-usage analyses support the DTU findings, the functional consequences of specific transcript isoforms remain to be experimentally validated. Third, longitudinal patient specimens spanning baseline, early on-treatment/MRD, and established resistance remain limited, constraining direct validation of the complete temporal trajectory in clinical samples. Finally, although our modeling prioritizes CSFs associated with MAPKi-induced transcript remodeling, direct perturbation studies will be required to determine whether these splicing factors regulate the identified isoform programs and influence drug tolerance. Future studies should determine whether the transient transcript-usage programs identified here can be detected in early on-treatment patient specimens and whether individual isoforms or their candidate regulators have functional roles in maintaining the drug-tolerant state or facilitating resistance evolution.

In summary, our findings support a model in which MAPKi treatment induces a broad transient splicing-associated transcript-usage state in surviving melanoma cells. These alterations have a much broader and different scope from those observed at the level of gene-level expression and are more predictive of the on-treatment adaptive phase than DEG-based features. They are enriched for resistance-associated signaling, metabolic, and developmental programs. By revealing adaptive pathway rewiring missed by conventional gene-expression analysis, transcript-level profiling provides a complementary framework for understanding melanoma plasticity and the evolution of acquired MAPKi resistance. The application of such analyses could potentially advance our understanding of the emergence of treatment resistance also in other indications and treatment regimes.

## Methods

### Datasets acquisition

Publicly available RNA-seq data from MAPKi-treated cell lines was downloaded from the GEO database with accession number GSE75299. This dataset has RNA-seq performed with four different melanoma cell lines, including untreated baseline, MAPKi on-treatment drug-tolerant cell lines, and resistance cells, comprising a total of 31 samples with multiple biological replicates at each time point. The dataset for the mouse PDX model of human melanoma was downloaded from GSE116237 and consisted of single-cell transcriptomic profiling of the PDX model at different stages: before treatment, two-time points during MAPKi treatment, and resistant to MAPKi. Both datasets capture critical time points, including the on-treatment phase, which is vital for understanding resistance mechanisms. Additionally, we identified the GSE75299 dataset, which includes pre-MAPKi and on-treatment melanoma patient biopsy samples, totaling 19 biopsies from 7 patients (7 no-treatment and 12 on-treatment). An independent murine melanoma bulk RNA-seq dataset (GSE103725) was also analyzed, comprising 19 samples across untreated (n = 4), on-treatment (n = 9), and resistant (n = 6) states.

### Generation of pseudo-bulk single-cell transcriptomes

To mitigate the sparse transcript-level detection inherent to single-cell RNA sequencing, which can limit the reliable quantification of individual transcript isoforms, we generated pseudo-bulk transcriptomes from the GSE116237 melanoma PDX single-cell RNA-seq dataset. For each of the four treatment phases (T0, Phase 1, Phase 2, and Phase 3), we randomly selected 50 cells as representative seed cells. For each seed cell, we identified its K-nearest neighbors within the same treatment phase based on Euclidean distances in the principal component analysis (PCA) space defined by the first ten principal components of gene-expression profiles. The neighborhood size (K) was set to 30% of the total number of cells within the corresponding treatment phase. Raw FASTQ files from cells belonging to each neighborhood were merged to generate a pseudo-bulk RNA-seq profile, which was subsequently used for transcript-level quantification and downstream analyses. By restricting neighborhood selection to cells from the same treatment phase, this approach increased transcript detection while preserving treatment-stage-specific expression characteristics. Because neighborhoods were constructed independently around each seed cell, individual cells could potentially contribute to multiple pseudo-bulk profiles. Accordingly, these computationally generated profiles represent aggregated transcriptomic measurements rather than independent biological replicates.

### Read mapping and Differential gene expression analysis

To map the sequence reads to the human genome build (GRCh38.p13), we utilized reference annotations obtained from the GENCODE release 43, and for the mouse, we used GENCODE release M36 (GRCm39). Transcript-level quantifications were performed using *salmon* (*29*). Following quantification, the ‘tximport ‘package (30) in R was employed to aggregate these transcript-level data into gene-level counts. This process involved importing the Salmon output, which includes estimates of transcript abundance and length, and then summarizing these to the gene level using a gene-to-transcript mapping. For the analysis of differentially expressed genes (DEGs), we utilized *DESeq2* (31) and followed the standard workflow designed for Salmon quantification files. Genes with a Benjamini-Hochberg adjusted *P value (padj)* < 0.05 were considered differentially expressed.

### Differential transcript usage (DTU) analysis

We used the R package DTUrtle (32) for the computation of differential transcript usage (DTU). To annotate the transcripts, we used the GENCODE release 43 genome annotation. The *run_drimseq* function was employed to calculate DTU genes for each group comparison. In order to enhance the statistical power of the analysis, we applied a filtering step where transcripts with low impact were excluded. Specifically, transcripts with less than 5% usage across all samples or a corresponding total gene expression of less than five counts for all samples were removed prior to statistical testing. Additionally, only genes that contained at least one high-impact transcript were taken into consideration. From this analysis, we obtained genes that exhibited an overall significant change in transcript usage, along with the corresponding transcripts driving the observed changes (both with an overall *FDR < 0.05*). Further, to reduce the number of analyzed transcripts per DTU gene, a post-hoc filtering step was implemented. Transcripts whose proportional expression deviated by less than 10% between samples were excluded. We considered only those transcripts that fulfilled the criterion that all samples within one group exhibited higher transcript usage compared to all samples in the other group.

### Differential exon usage analysis

‘Percent spliced in ’or the PSI value is a commonly used metric to quantify the inclusion level of exons from short-reads RNA-seq data. We used rMATs (33) to compute the PSI value of each alternative splicing event defined based on the annotations in the GTF file from Gencode V43. We first ran the STAR two-pass alignment (34) to reliably map the short-reads from raw .fastq files to splicing junctions and supplied the resulting transcriptomic bam files to rMATs to quantify the inclusion level of five different kinds of events including skip exons (SE), alternative 5 ’and 3 ’splice sites(A5SS and A3SS), mutually exclusive exons (MXE) and intron retention (IR). We ran rMATs separately for each input sample and the resulting .psi files were merged across all samples to create a matrix (event on by sample) of PSI values using custom scripts in R. As an alternative, we also used SUPPA2 (35) to quantify the exon inclusion based on the precomputed transcript level TPM values from Salmon.

To validate the transiently upregulated transcripts (TUTs) using differential exon usages, we first identified the splicing events that were part of the TUTs. We extracted this mapping information between transcripts and splicing events from the .ioe created by SUPPA2. We used the .ioe file to map splicing events quantified by both SUPPA2 and rMATs. Since rMATs used different genomic coordinate conventions to designate splicing events, we created a custom R script to translate the coordinates of splicing events from rMATs to SUPPA2. After identifying the splicing events mapping to TUTs, we modeled the cell line labels of the MAPKi treated data (i.e baseline and on-treatment, coded as 0 and 1 respectively) as response variable and PSI value of each alternative splicing event as predictors. Since multiple cell lines were used for experiments in GSE75299, we used a mixed-effects regression model to account for cross-cell line variability. For this purpose, we used lme4 package in R by specifying binomial as link function. A positive regression coefficient for any given exon would indicate that its usage in on-treatment samples is relatively higher as compared to the untreated baseline.

### Machine Learning-Based Stratification of No-Treatment and On-Treatment Phases Using DTUs and DEGs

We employed machine-learning classifiers to compare the ability of transcript-usage and gene-expression features to distinguish no-treatment from on-treatment samples. Two separate feature matrices were generated: one containing the usage values of DTU transcripts and the other containing the expression values of DEGs. Classification models were first trained using all available DTU or DEG features. We then performed feature selection independently for the DTU and DEG datasets using a linear support vector classifier (36) with L1 regularization (SVC-L1). Within each feature set, the SVC-L1 model assigns a coefficient to every transcript or gene; features with coefficients reduced to zero were discarded, whereas features with non-zero coefficients were retained as the selected DTU or DEG features. These independently selected feature subsets were subsequently used to train reduced binary classification models for distinguishing no-treatment from on-treatment samples.

### Pathway overrepresentation analysis and enrichment score calculation

Based on the literature survey, we first selected pathways that are crucial for MAPKi resistance in melanoma. Then, we extracted the genes associated with each pathway from the Reactome database (37). For each pathway, an enrichment score was calculated to quantify the degree to which sets of differential transcript usage (DTU) genes and differentially expressed genes (DEG) were overrepresented. The enrichment score was defined as the ratio of the observed number of genes in a given pathway to the expected number based on the pathway’s representation in the background gene set. Specifically, for a given set of genes G (either DTU or DEG) and a pathway P, the observed count O_G was the number of genes in G that were also in P. The expected count E_G was calculated as the product of the size of G, and the proportion of P in the background gene set B, i.e., E_G = |G| * (|P| / |B|), where |·| denotes set size. The enrichment score (ES) was then ES = O_G / E_G. Enrichment scores for each pathway were calculated for both DTU and DEG gene sets.

To assess the statistical significance of the observed enrichment, Fisher’s Exact Test was employed, comparing the observed overlap to what would be expected by chance, given the sizes of the gene sets and the background. The background gene set comprised all genes mapped in the RNA-seq data, ensuring the calculation of enrichment scores and p-values was contextualized within the scope of the experimental data.

### ssGSEA analysis of DTUs and DEGs

To investigate the association of differential transcript usage (DTU) and differential gene expression (DEG) with MAPKi resistance-associated signaling programs, we performed single-sample gene set enrichment analysis (ssGSEA) using the GSVA package (38) in R. For each pathway of interest, enrichment scores were calculated independently for transcript-usage and gene-expression data across samples representing no-treatment, on-treatment, and acquired-resistant states. ssGSEA was performed using the rank-based enrichment approach implemented in GSVA, generating sample-specific enrichment scores for the selected pathways. These scores were subsequently compared across treatment states to characterize dynamic changes in MAPKi resistance-associated signaling programs and to evaluate the complementary information captured by transcript-level and gene-level analyses.

### Comparison of splicing observed in on-treatment melanoma cells with developmental splicing and functional enrichment analysis

To investigate the developmental roots of splicing changes underlying the transient response to MAPKi, we used an RNA-seq dataset from a previous study where pluripotent stem cells were differentiated into melanoblasts and then to melanocytes. This study consisted of multiple experiments where certain tumor suppressor genes were knocked down, followed by doxycycline treatment to induce BRAFV600E. We only focused on wildtype and untreated experiments from this study to focus on the splicing events specific to normal developing melanoblasts and melanocytes. Using three biological replicates for melanoblasts and melanocytes from wildtype and untreated experiments, we first computed the transcript usage in each sample using SUPPA2 and identified the differential splicing events between these two conditions using a Wilcoxon’s rank sum test followed by the multiple testing correction of the resulting p-values. Transcripts with absolute delta PSI > 0.1 and FDR < 0.05 were termed as differential between melanocytes and melanoblasts. We intersected these differential transcripts with the list of TUTs and TDTs and made a boxplot of the delta PSI (melanoblasts - melanocytes) values.

### Model for regulation of splicing events and identification of critical splicing factors

To identify the potential splicing regulators of the TUTs, we shortlisted a total of 442 genes with potential splicing regulatory functions from the Amigo database. We first quantile normalized the gene expression data from all the MAPKi treated and untreated cell lines from GSE75299, and using the gene expression of these 442 genes, we used a partial least square regression (PLSR) approach to model the mean inclusion level of TUTs. The PLSR approach can effectively handle the multi-collinearity among the predictor variables while maximizing the covariance between the response and predictors. We used the PLS package in R for this purpose. We evaluated the significance of PLSR regression coefficients through jackknife resampling which computes a t-statistics by iteratively refitting the PLSR model by dropping one observation at a time. Splicing factors with positive regression coefficient in the first component and FDR corrected p-value < 0.05 were termed as CSFs.

---

**Supplementary Figure 1.**
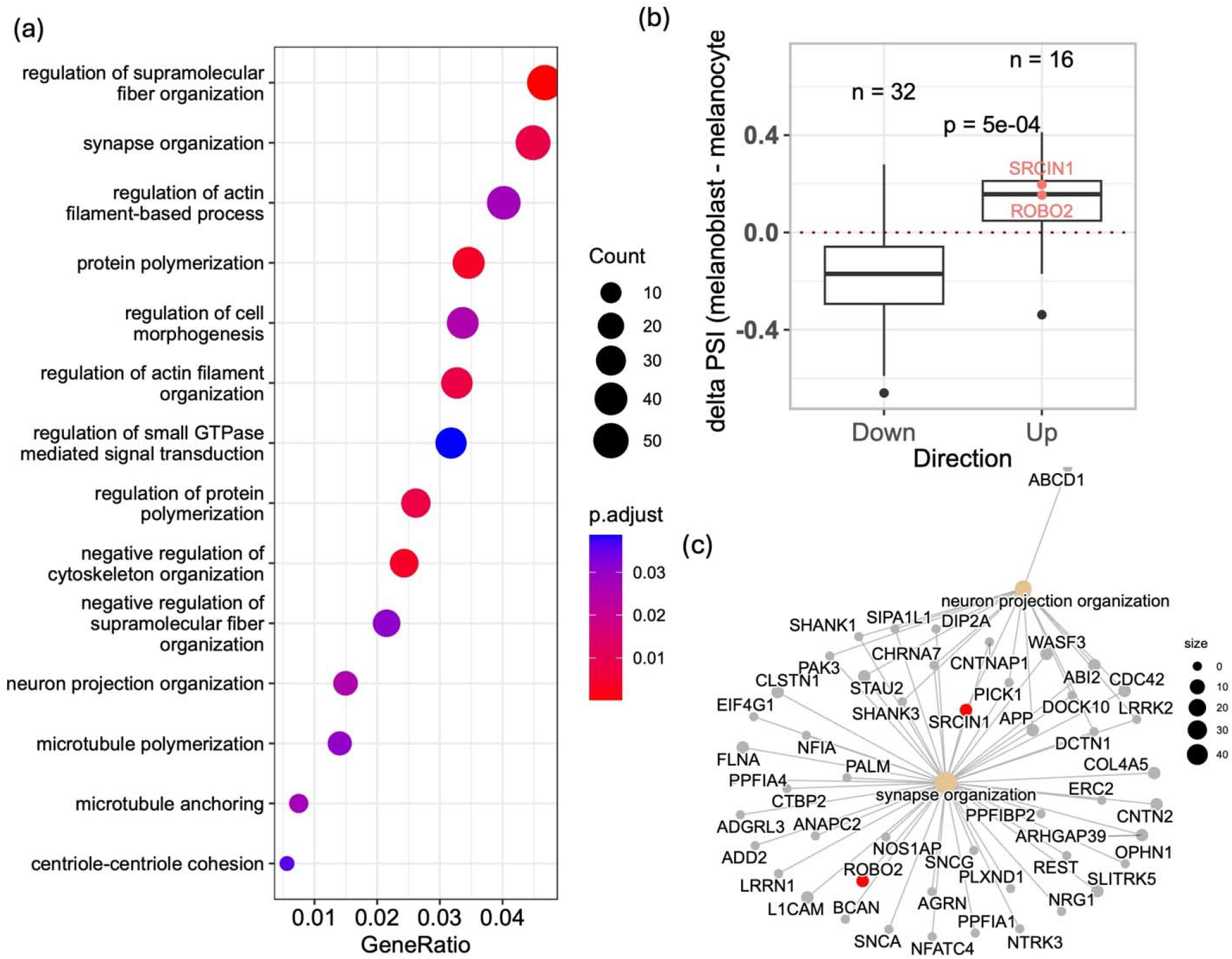
Developmental splicing programs associated with melanoblast differentiation and MAPKi-induced transcript remodeling. (a) Gene Ontology enrichment analysis of genes associated with transcripts showing higher inclusion in melanoblasts relative to differentiated melanocytes. Enriched biological processes were dominated by cytoskeletal and cellular-organization programs, including regulation of actin filament organization, supramolecular fiber organization, cell morphogenesis, protein polymerization, and neuron projection organization. Dot size represents the number of genes associated with each term, and color indicates the *adjusted P value*. These enrichments are consistent with the migratory and spindle-shaped phenotype of melanoblast precursor cells. (b) Comparison of melanoblast-versus-melanocyte differential transcript usage for MAPKi-responsive transcripts using a more stringent set of developmental splicing events (absolute ΔPSI> 0.1 and *FDR < 0.05*). Transcripts increased during the MAPKi on-treatment state showed significantly higher melanoblast-associated ΔPSI values than transcripts decreased during treatment (*P = 5 × 10*⁻⁴; n = 16 upregulated and n = 32 downregulated transcripts), further supporting preferential reactivation of melanoblast-associated isoform usage during MAPKi adaptation. Representative transcripts including SRCIN1 and ROBO2 are highlighted. (c)Gene-pathway network illustrating genes contributing to the enriched synapse organization program identified among melanoblast-associated transcripts. The central node represents the enriched biological process, with connected nodes denoting contributing genes; ROBO2 is highlighted as a representative developmentally regulated transcript associated with this program.

## Data availability

All transcriptomic datasets analyzed in this study are publicly available from the NCBI Gene Expression Omnibus (GEO), including GSE75299 (MAPKi-treated melanoma cell lines and patient samples), GSE103725 (murine melanoma model), and GSE116237 (melanoma patient-derived xenograft single-cell RNA-seq). The developmental melanoblast and melanocyte RNA-seq data were obtained from the study by Baggiolini et al. (2021).

## Acknowledgements

This research is supported in part by the Intramural Research Program of the National Institutes of Health, National Cancer Institute, Center for Cancer Research. This work has utilized the computational resources of the NIH HPC Biowulf cluster (http://hpc.nih.gov). ChatGPT Plus was used strictly to refine certain existing paragraphs to improve sentence constructions.

