## Supplementary figures and images for "Transient Splicing-Associated Isoform Remodeling Defines Early MAPKi Resistance Adaptation in Melanoma"

### Supplementary Figure 1

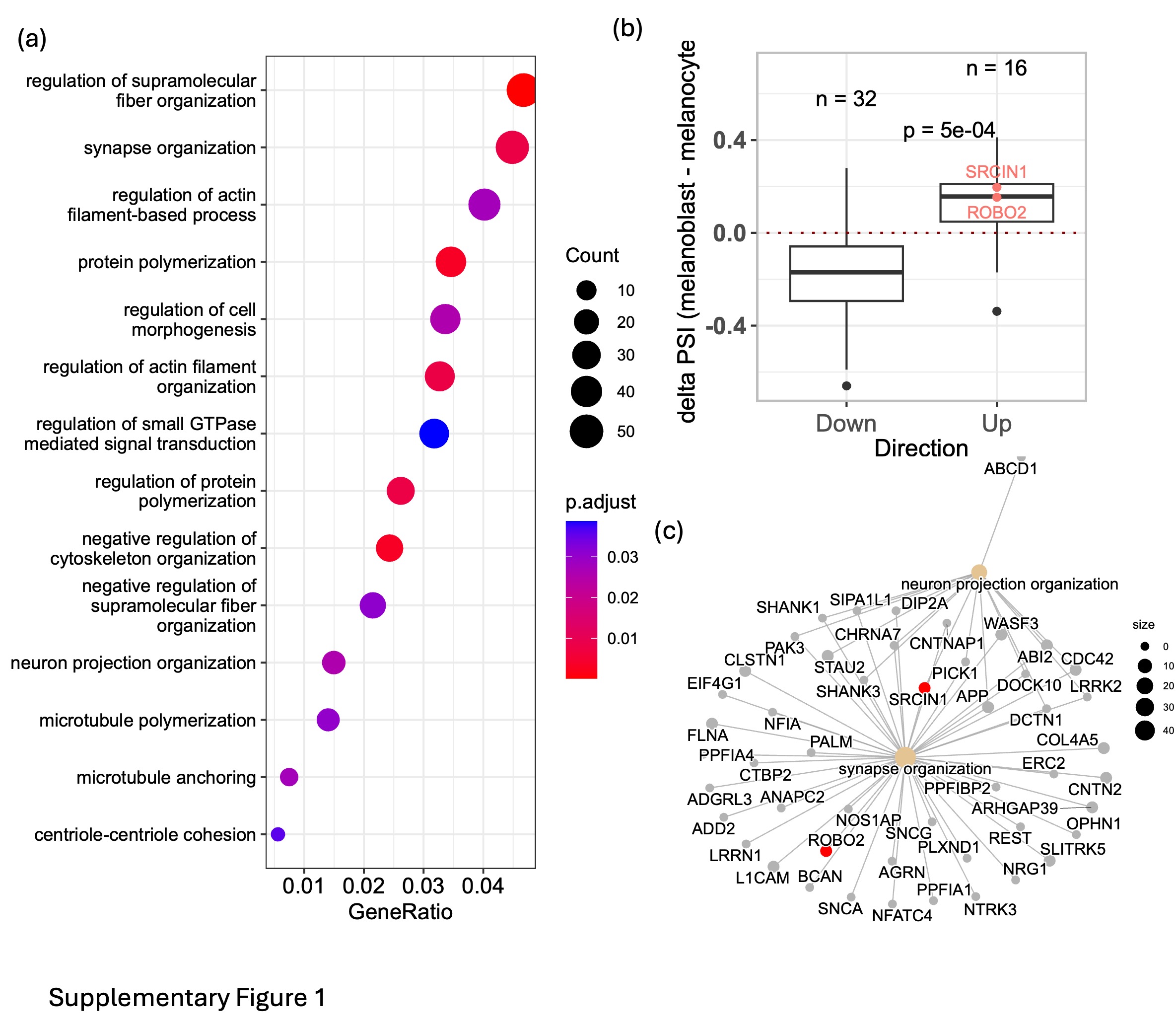
